# Distinct representations of an anxiogenic environment in different cell types of the ventral hippocampus

**DOI:** 10.1101/2025.03.07.641976

**Authors:** Jack E Berry, Jessica C Jimenez, Kenechukwu Michael Ogbu, Nicholas Manfred, Karly Tegang, Stefano Fusi, René Hen, Clay Lacefield, Wei-Li Chang

## Abstract

In addition to its role in episodic memory and spatial navigation, the hippocampus has also been found to influence mood-related disorders such as anxiety and depression. These seemingly distinct roles are consistent with a functional dissociation between the two anatomical poles of the hippocampus: whereas the dorsal portion of the hippocampus in rodents is necessary for spatial tasks, the ventral portion controls affective behaviors. We have recently found that neurons in the ventral, but not dorsal, CA1 area of mice encode anxiety-related information (i.e. are “anxiety cells”) in diverse defensive and exploratory behaviors. Still it is unclear how general threat-related information is computed within the hippocampal circuit. In this work, we have examined how distinct hippocampal subregions and cell types encode anxiety-related information by imaging calcium activity in large populations of genetically-defined neurons in the ventral hippocampus while mice explore the elevated plus maze (EPM), a conflict-based anxiety test. We compared the neural encoding of task-related features within the ventral CA1 (vCA1) and ventral dentate gyrus (vDG) regions in order to examine the emergence of anxiety-related activity through the hippocampal circuit. We found that granule cells (vGCs) of the vDG represented similar valence information to neurons in vCA1 in the form of arm type-specific encoding in the EPM, which suggests that encoding of anxiety-related features is already present at this first stage of hippocampal processing. When compared with ventral granule cells (vGCs), ventral mossy cells (vMCs) underlying the DG had stronger spatial encoding and less valence encoding, suggesting that they may be more functionally connected with the highly spatially sensitive dorsal hippocampus. Together these findings will help to understand the encoding of anxiety-related information in the hippocampus and how it relates to neural circuit defects in mood-related disorders.

## INTRODUCTION

The ventral portion of the hippocampus (vHPC) has been implicated in emotional behaviors in humans^1^, non-human primates, and rodents^2^. Both ventral dentate gyrus (vDG)^3,4^ and ventral CA1^5,6^ areas contribute to natural avoidance behavior in approach-avoidance tasks, but it is unclear what aspects of such behaviors are encoded by these regions. Recent studies have found that pyramidal neurons in ventral, but not dorsal, CA1 exhibit anxiety-related neural activity in the elevated plus maze (EPM)^5,6^. Furthermore, vCA1 anxiety-related neurons are more likely to project to the medial prefrontal cortex and lateral hypothalamus than other brain areas such as amygdala and nucleus accumbens, which suggests involvement in specific downstream circuits and behaviors. While these previous studies identified single-cell properties of vCA1 neurons in the EPM, it is still unclear how these neurons compute anxiety-related information, and whether valence is encoded upstream of vCA1 in the hippocampal trisynaptic circuit.

Here, we recorded ventral hippocampus calcium signals with a miniature head-mounted microscope while mice performed a classic approach-avoidance task, the elevated plus maze (EPM), to assess how anxiety-related information is encoded in vDG, the initial stage of hippocampal processing, as well as in vCA1 (**Figure 1A-C, S1**). Furthermore, the vDG is unique within the hippocampus in that it consists of two distinct subpopulations of excitatory neurons: granule cells (GCs), the principal projection neurons, and mossy cells (MCs) which lie in the hilar region underlying the DG and send feedback projections to GCs and inhibitory interneurons. These cell types differ in their intrinsic physiology, neuromodulation, and local anatomical connectivity^7^; therefore, it is possible that they are responsible for representing distinct aspects of an anxiogenic context. We used Dock10-Cre and Drd2-Cre mouse lines to express Cre-dependent GCaMP AAV selectively in vGCs and vMCs, respectively, in order to record calcium activity in these distinct vDG cell types during EPM exploration (**Figure 1D-E**). Additionally, calcium imaging data acquired previously^5^ from vCA1 cells in the EPM were re-analyzed in the same analysis pipeline to determine to what extent vDG and vCA1 encode anxiety-related information in a similar manner (**Figure 1F**).

**Figure 1.**
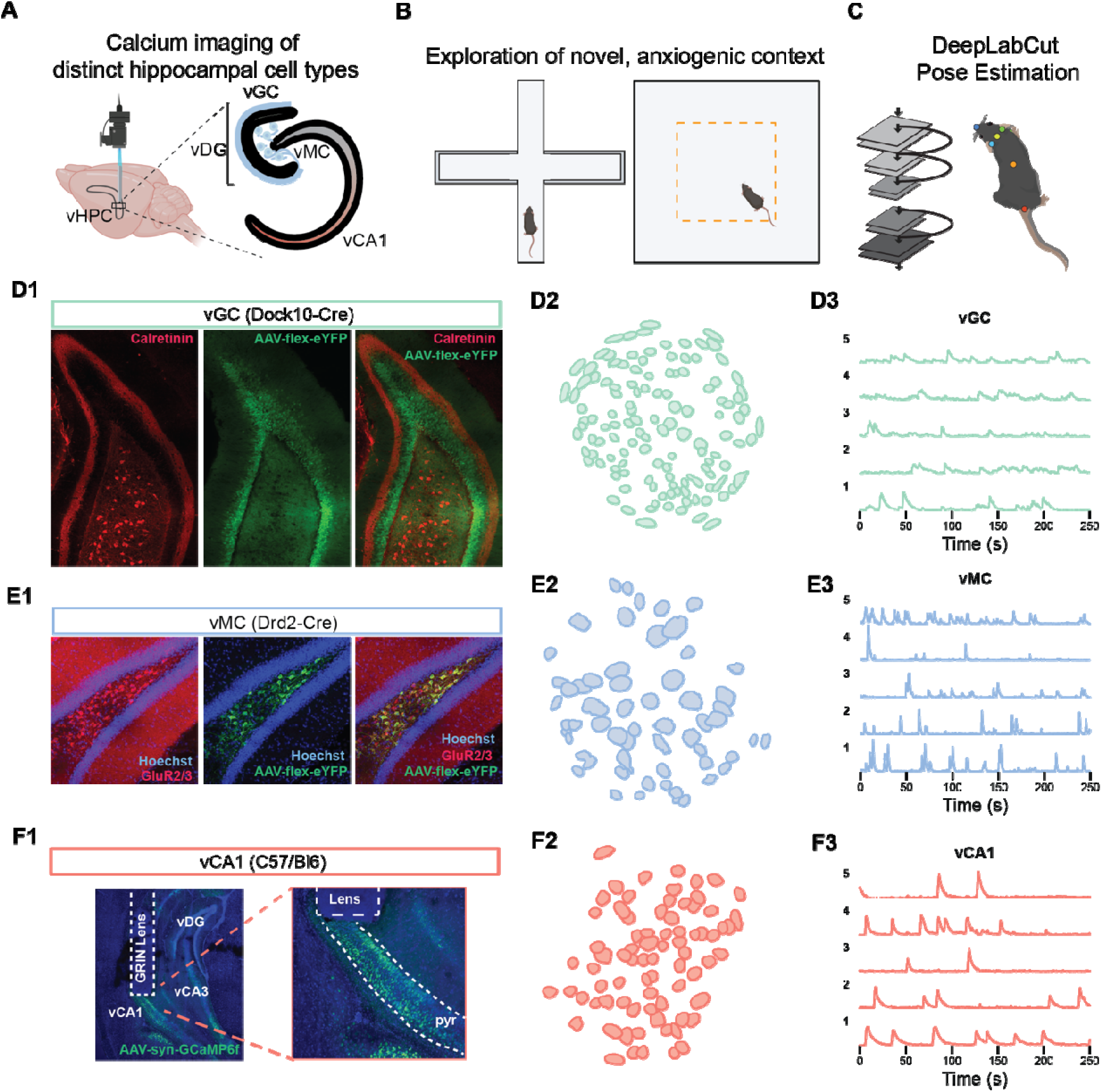
Calcium imaging of distinct ventral hippocampal cell types during exploration of a novel, anxiogenic environment. (A) Schematic of experimental design. GCaMP-expressing AAV was injected into either vDG or vCA1, and a GRIN lens was then inserted above the region to provide optical access.(B) Calcium activity was recorded while mice explored a novel elevated plus maze and open field. (C) Using the DeepLabCut software package, a neural network was trained on a manually labeled set of images to recognize six body parts: nose, left ear, right ear, head, body, and tail base. The location of the head was used throughout this study to determine the location of the mouse. (D) vGC calcium imaging. (D1) The Dock10-Cre line was used to target vGCs. (D2) Representative field of view depicting cell contours of segmented vGCs. (D3) Representative Ca^2+^ traces from five vGCs. (E) vMC Ca^2+^ imaging. (E1) The Drd2-Cre line was used to target vMCs. (E2) Representative field of view depicting cell contours of segmented vMCs. (E3) Representative Ca^2+^ traces from five vMCs. (F) vCA1 Ca^2+^ imaging. (F1) Ca^2+^ videos from Jimenez et al., 2018 were reanalyzed in the same analysis pipeline as the vGC and vMC videos. (F2) Representative field of view depicting cell contours of segmented vCA1 neurons. (E3) Representative Ca^2+^ traces from five vCA1 neurons.

## RESULTS

### Ventral hippocampal neurons exhibit increased activity during open arm exploration, head dips, and stretching

We recorded calcium activity from populations of vGC, vMC, and vCA1 neurons while the mouse freely explored a novel EPM for ten minutes. Consistent with previous reports^5,6^, vCA1 neurons exhibited increased activity in the anxiogenic open arms and during head dips, which are associated with risk assessment^8^ (**Figure 2A**). Despite the average increase in activity in the open arms, the population consisted of a mixture of neurons that were selectively active in each of the three zones, open arms, closed arms, and the center region (**Figure 2B**). Neurons were classified as selective for a particular zone by comparing the calcium rate increase to a shuffled distribution, as previously described^5^ (see Methods). Whereas 37% of vCA1 neurons were selective for the open arms, only 17% were selective for the closed arms. vGCs and vMCs were enriched for open arm-selective cells, with a greater proportion of vMCs (48%) preferring open arms than either vGCs (31%) or vCA1 (37%) (**Figure 2B**). The degree to which the population activity increased in the open arms was correlated with the anxiety level of the mouse, as measured by a decrease in the time spent in the open arms (**Figure S2**).

**Figure 2.**
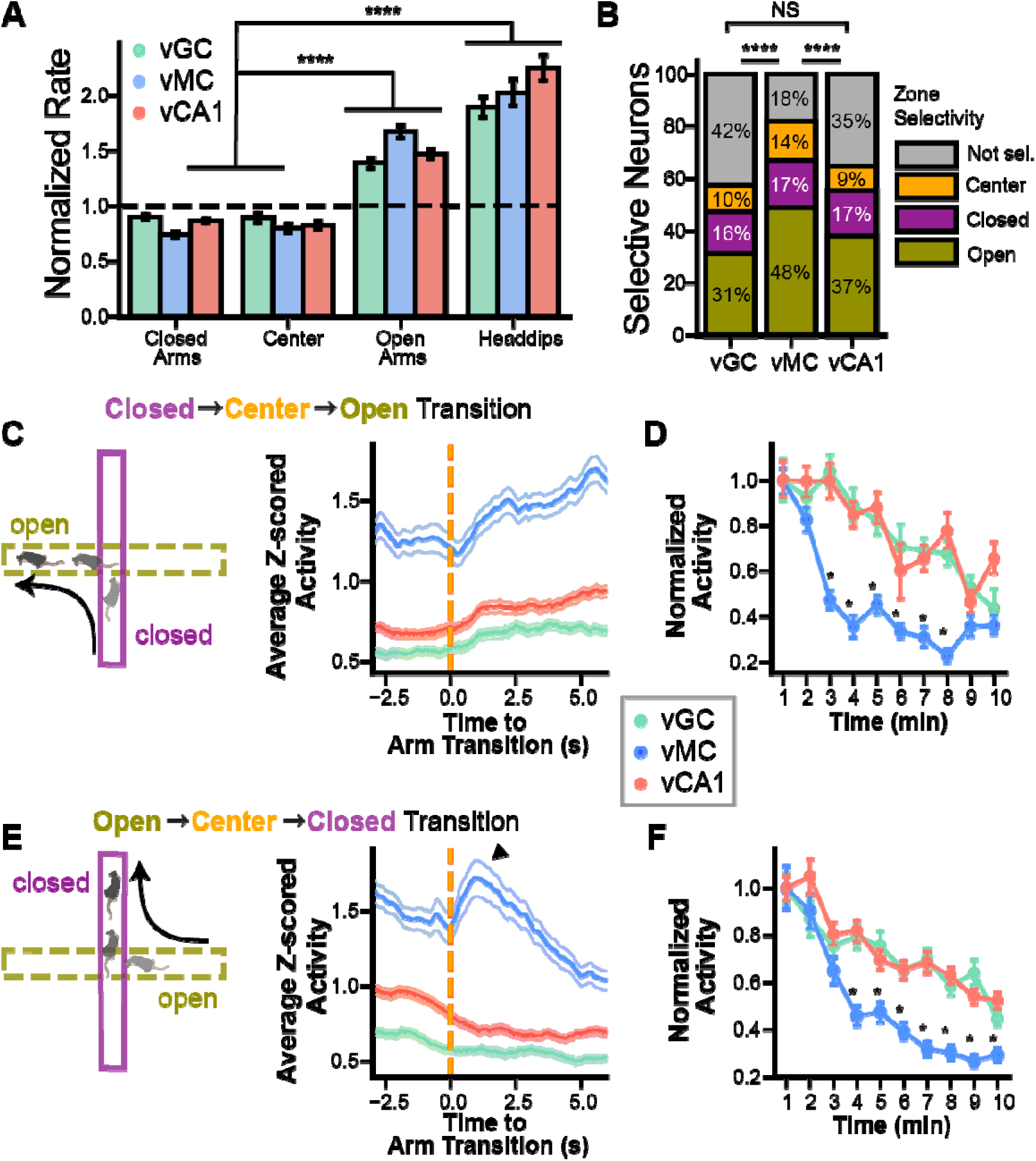
vMCs are strongly modulated by novelty, and all vHPC cell types exhibit increased activity during risk assessment in the EPM. (A) Normalized Ca^2+^ activity rate in the closed arms, center, and open arms, and during head dips and stretching behaviors. Normalized rates are computed for each neuron by dividing the mean rate during the behavioral epoch by the mean rate during the entire session. In all three cell types, rates are significantly elevated when mice explore the open arms, and during head dipping and stretching episodes (p<0.0001 for all groups, Mann Whitney U). (B) All three vHPC cell populations are enriched in cells that are selectively active in the open arms. Cell selectivit was assessed by comparing the rate difference between in- and out- of zone activity with a shuffle distribution, as previously described(Jimenez et al., 2018). (C) vHPC activity increases as the mouse transitions from closed to open arms. (Left) Cartoon of mouse leaving closed and entering open arms. (Right) Mean activity for each cell type triggered on entering an open arm following a <2s visit to the center. (D) Ca^2+^ activity in the open arms decreases with exposure, but more dramatically in vMCs. (E) vHPC activity decreases as the mouse transitions from open to closed arms. (Left) Cartoon of mouse leaving open and entering closed arm. (Right) Activity of each cell type during triggered on entering the closed arm. Note the transient increase in vMC activity upon closed arm entry before declining. (F) Averaged rates per minute, normalized to the first minute of EPM exploration. As with the open arms, Ca^2+^ activity in the closed arms also decreases with exposure, however again vMCs exhibit a significantly faster decline in rate compared to vGCs and vCA1. Stars indicate significant (p<0.05; Kruskal-Wallis) differences between cell groups. Error bars indicate 68% bootstrap confidence interval to approximate SEM. N_vGC,mice_ = 5; N_vMC,mice_ = 8; N_vCA1,mice_=6. N_vGC,cells_ = 391; N_vMC,cells_ = 314, N_vCA1,cells_ = 534.

### vMCs encode contextual novelty

In addition to being a generally anxiogenic environment, the animals are also exposed to the EPM as a novel context. Stimulus novelty has important influence on mnemonic function, exploratory behavior, and hippocampal activity. Furthermore, novelty detection has been proposed as a potential role for mossy cells in particular^7^. For a neuron to encode novelty, its activity should be higher during the initial exposures to each arm than at the end of the session (**Figure 2D,F, S2**). To evaluate the rate at which calcium activity changed during exposure to the novel arena, we normalized each neuron’s calcium event rates over time to its rate during the first minute. While vMCs showed higher event rates than vGCs or vCA1 neurons in the EPM, they also notably exhibited faster habituation, with the normalized mean rate decreasing by 50% after only 2 minutes in the open arms, and 3 minutes in the closed arms (**Figure 2D,F, S2**). In comparison, vGCs and vCA1 neurons did not return to similar levels until the last 2 minutes of the 10-minute session, showing that vMCs had a stronger response to the novelty of the EPM arms.

To further explore the role of vHPC neuron subtypes in encoding novelty, we examined neural activity triggered on arm transitions. In all three cell types, vCA1, vGCs and vMCs, the average calcium signals showed an increase upon entering an open arm, and a decrease upon entering a closed arm (**Figure 2C,E**), which is consistent with the higher average event rates in open arms from the previous cell selectivity analysis. However, vMCs were unique in that they had a brief increase in activity upon entering a closed arm, before declining below pre-entry levels (**Figure 2E**, black arrowhead). This activity pattern is consistent with the idea that vMCs are particularly sensitive to novel stimuli or changes in context.

### vMCs robustly encode spatial information

In addition to its use as a test of approach-avoidance behavior, the EPM also involves spatial exploration of the complex maze geometry by the mouse. We computed the significance of spatial information (SSI)^9,10^ to evaluate the degree to which individual neurons encode spatial information in the EPM. In agreement with early studies^11^, vGCs and vCA1 neurons had low spatial information, with a mean SSI of 1.46 and 1.52, respectively. However, vMCs had a mean SSI of 3.69, which is significantly higher than vGCs or vCA1 (**Figure 3A-E**), suggesting these neurons may perform a distinct function within the ventral hippocampus that involves more precise spatial encoding.

**Figure 3.**
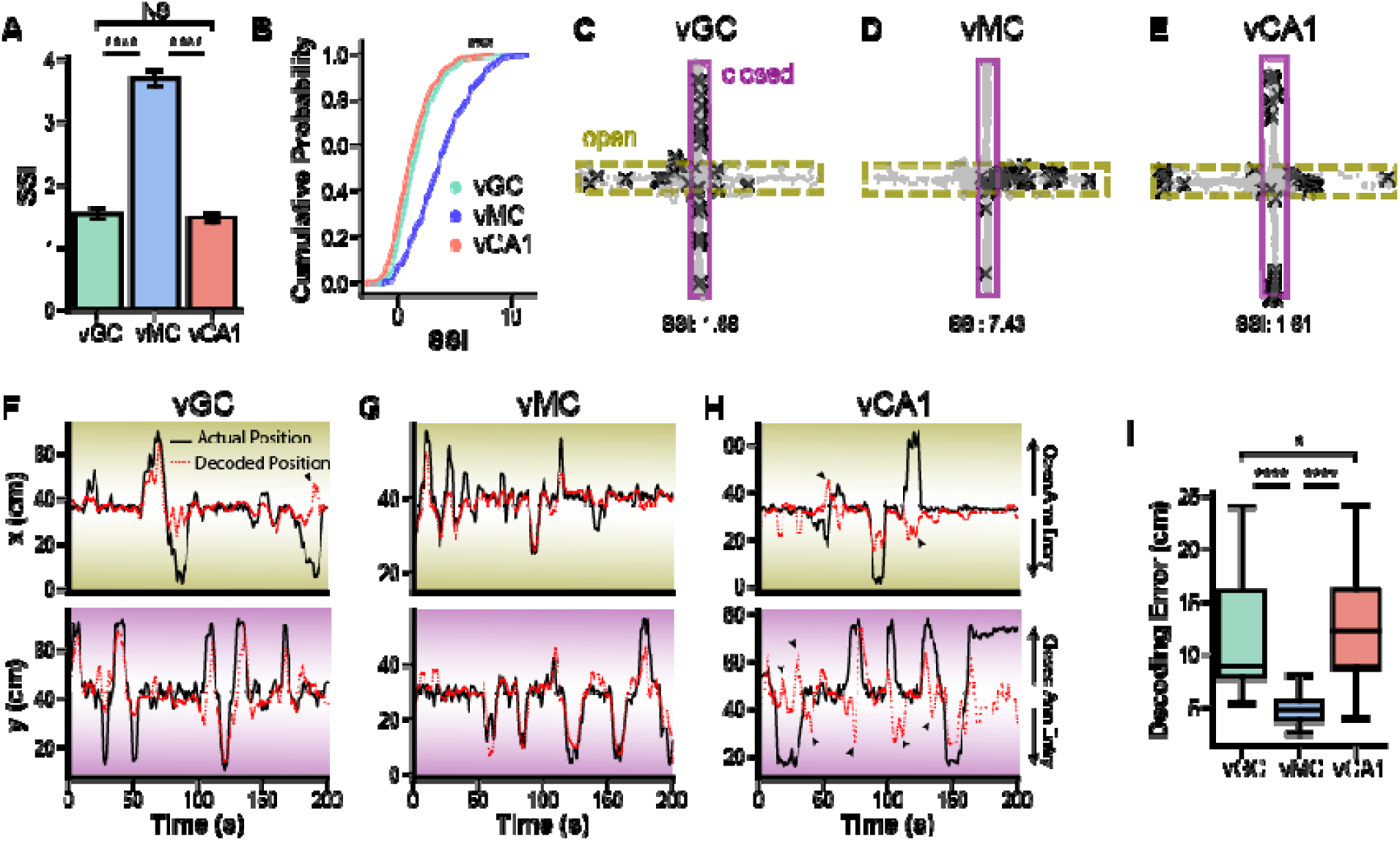
vMCs, but not vGCs or vCA1, robustly encode spatial position in a novel environment. (A and B) Standardized spatial information (SSI) is higher in vMCs than vGCs or vCA1. (p < 0.0001; Mann-Whitney U, N_vGC,cells_ = 391; N_vMC,cells_ = 314; N_vCA1,cells_=534), whereas there was no difference in SSI between vGCs and vCA1 (p = 0.19; Mann-Whitney U) (C-E) Representative spatial activity patterns for vGC (C), vMC (D) and vCA1 (E) neurons. (F-I) The spatial information content encoded by the neural population was assessed by decoding the position of the mouse using neural activity. The arena wa divided into a grid of 16×16 squares roughly 4×4cm. A support vector classifier with a linear kernel was trained to predict the spatial bin of the mouse using 10-fold validation. (F-H) Example decoder performance for three mice using vGCs (F), vMCs (G) or vCA1 (H). The x (top) and y (bottom) position are shown with actual position in black, and decoded position in red. Note that due to the spatial configuration of the EPM, position deviations in the x direction correspond to open arm entries, wherea deviations in the y direction correspond to closed arm entries. Position decoding errors in vGC and vCA1 mice often occur due to predicting the mouse has entered the wrong arm of the same valence (arrowheads). (I) The decoding error, measured as the distance between decoded and actual position, is significantly lower in vMCs than in either vGC or vCA1 (p < 0.0001; Kruskal-Wallis; post-hoc Mann-Whitney U: p_vGC,vMC_ <0.0001, v_vCA1,vMC_ < 0.0001, p_vGC,vCA1_ = 0.019; N_vGC,mice_ = 5; N_vMC,mice_ = 8; N_vCA1,mice_=6).

To further investigate the spatial encoding properties of vHPC neurons, we used a support vector classifier to decode the mouse’s position, as previously described for dorsal GCs and CA1^10^. We assessed the significance of the decoder performance by comparing the decoding error (distance between decoded and actual position, **Figure 3F-H**) to that of a decoder trained on shuffled position data (see Methods). We were able to decode the position of the mouse from all three cell types, however performance was significantly better when using vMCs (**Figure 3I, S4**). Notably, decoding errors in vGCs and vCA1 were often due to predicting an entry into the wrong arm of the same valence (e.g. **Figure 3H**, arrowheads). This is consistent with vGC and vCA1 neurons encoding a more general concept of an arm of a particular valence, rather than position.

Spatial information is not typically examined in the EPM due to the maze’s irregular shape and the mouse’s uneven occupancy of the arena, both of which can affect traditional spatial information measures. To confirm our findings of spatial encoding in distinct vHPC populations, we recorded calcium activity from these neurons as the mouse explored an open field (**Figure S5**). As in the EPM, vGCs and vCA1 neurons had low spatial information in the open field, whereas vMCs had significantly higher SSI. Furthermore, a linear decoder was able to successfully decode position in the open field from all three cell types, but performance was significantly higher in the vMC group.

### vGCs and vCA1 encode arm type

We next investigated whether distinct vHPC neuron subpopulations encode information about arm type in the EPM, and thereby underlie valence encoding in anxiety cells. The EPM consists of four spatially distinct arms that can be grouped in pairs based on the type or valence of the arm, where open arms are thought to be anxiogenic while closed arms are comparatively safe. We found that the three vHPC populations, vGCs, vMCs, and vCA1 neurons, each exhibited increased average activity during open arm exploration. However it is unclear whether neurons are truly encoding the negative valence of the open arms as a general feature, or rather have more cells that encode each open arm independently. To examine this possibility, we first computed the EPM score, a single-cell measure of arm type coding based on the ratio of activity differences between same vs. different arm types^6,12^. There was a small difference in the EPM scores of vGCs, vMCs, and vCA1 neurons (Anderson k-samples test, p<0.05), with a left-shift in the distribution of EPM score among vMCs compared with vCA1 and vGCs (**Figure 4A,B**; p<0.05, post-hoc Anderson-Darling test with Bonferroni correction), suggesting weaker arm/valence encoding of individual vMC neurons.

**Figure 4.**
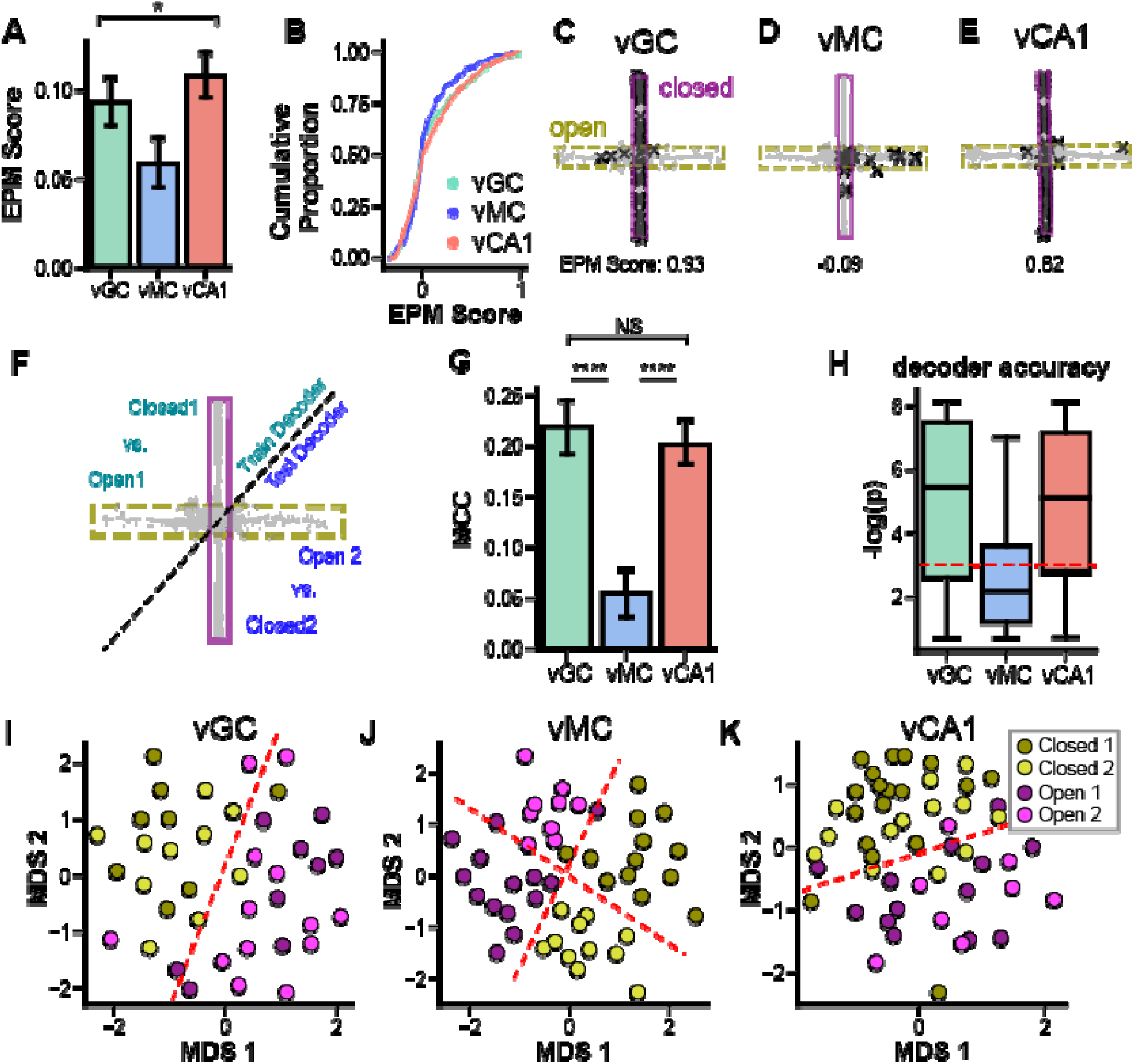
vGCs and vCA1, but not vMCs, encode arm type. (A and B) Across the entire population, EPM score is lower in vMCs than either vGC or vCA1. (p = 0.014; k-sample Anderson-Darling, N_vGC,cells_ = 391; N_vMC,cells_ = 314; N_vCA1,cells_=534), (C-E) Example activity patterns of neurons with high (C,E) and low EPM score. Note that EPM score can be high for either closed- or open-preferring neurons. (F-H) A linear decoder was used to ascertain whether the geometry of the neural representations allows for generalization to other contexts with similar valence. (F) Schematic of decoding strategy. A SVC with linear kernel was trained to distinguish a single pair of different-valence arms (e.g. Open1 vs Closed1) and tested on distinguishing the remaining pair of arms (i.e. Open2 vs Closed2). This process was completed for all 4 possible arm pairings, and the performance the 4 decoders was averaged to determine overall performance. (G) Decoder performance was measured with the Matthews Correlation Coefficient (MCC), an unbiased measure of accuracy that ranges from -1 to 1, with 0 indicating chance-level performance. Arm valence was generalizable using vGCs and vCA1, but not vMCs (p < 0.001; Kruskal-Wallis; ad-hoc Mann Whitney U: p_vGC,vCA1_ = 0.4, p_vGC,vMC_ < 0.0001; p_vCA1,vMC_ < 0.0001) (H) Significance of decoder accuracy measured as –log(p-value). P-value was computed using a Mann Whitney test to determine whether the true performance was significantly different from decoding on shuffled data. Red dotted line indicates –log(0.05). (I-K) The geometry of the neural representation of EPM arm valence was further investigated by constructing multidimensional scaling (MDS) plots. Each circle represents a population vector for each entry into Open1 (purple), Open2 (magenta), Closed1 (olive green), or Closed2 (lime green) EPM arms. Shown here are representative MDS plots for vGC (I), vMC (J), and vCA1 (K). vMCs represent each arm as a distinct context, whereas vGCs and vCA1 neurons group arms of similar valence (i.e. open and closed). Red dotted lines depict a hypothetical linear classifier trained in similar manner as in (F).

We next assessed whether these three cell types encode information about arm type at the population level using a linear decoder. We trained the decoder to distinguish open and closed arms using training data from a pair of opposite-type arms (e.g. Open1 and Closed1), and then tested whether this decoder accurately classifies the arm type of the mouse using data from when the mouse explored the remaining arms (i.e. Open2 and Closed2, **Figure 4F**). By restricting the training data to one arm of each type, this strategy tests whether the population codes arm type in an abstract manner, which enables generalization^13^. Arm type was encoded by both vGCs and vCA1, indicating that these populations represent this feature of the EPM in an abstract manner, including potentially their behavioral relevance or valence. Conversely, arm type was not able to be decoded from vMCs (**Figure 4G,H**), which is consistent with the finding that these cells have lower EPM score but higher spatial tuning.

To visualize the distinct coding strategies for these vHPC populations, a population vector was constructed for each arm entry and represented in two dimensions using multidimensional scaling (MDS). The MDS plots (**Figure 4I-K**) demonstrate that whereas the vMC population encodes each arm as a distinct context, vGCs and vCA1 populations cluster each open and closed arm together, therefore representing arm type in an abstract fashion that may underlie the valence encoding properties of anxiety cells in the ventral hippocampus.

### vGCs, but not vMCs, are necessary for avoidance in the EPM

Given the differences we have observed in the encoding of arm valence in vDG neuron subtypes, we hypothesized that vGCs, but not vMCs would be important for natural avoidance behavior in the EPM. To test this possibility, we injected a Cre-dependent AAV expressing either inhibitory DREADD (AAV-DIO-hM4Di-mCherry) or control AAV (AAV-DIO-mCherry) in the vDG of either Dock10-Cre, or Drd2-Cre mice to selectively inhibit vGCs or vMCs, respectively (**Figure 5A,E**). Chemogenetic inhibition of vGCs increased the total distance traveled, time spent in the open arms, and percent of distance traveled in open arms, consistent with a role for vGC “anxiety cells” in driving avoidance behavior in the EPM (**Figure 5B-D**). However, silencing vMCs failed to affect each of these parameters (**Figure 5F-H**), indicating that vGCs are critical for approach-avoidance behavior, while vMCs are not.

**Figure 5.**
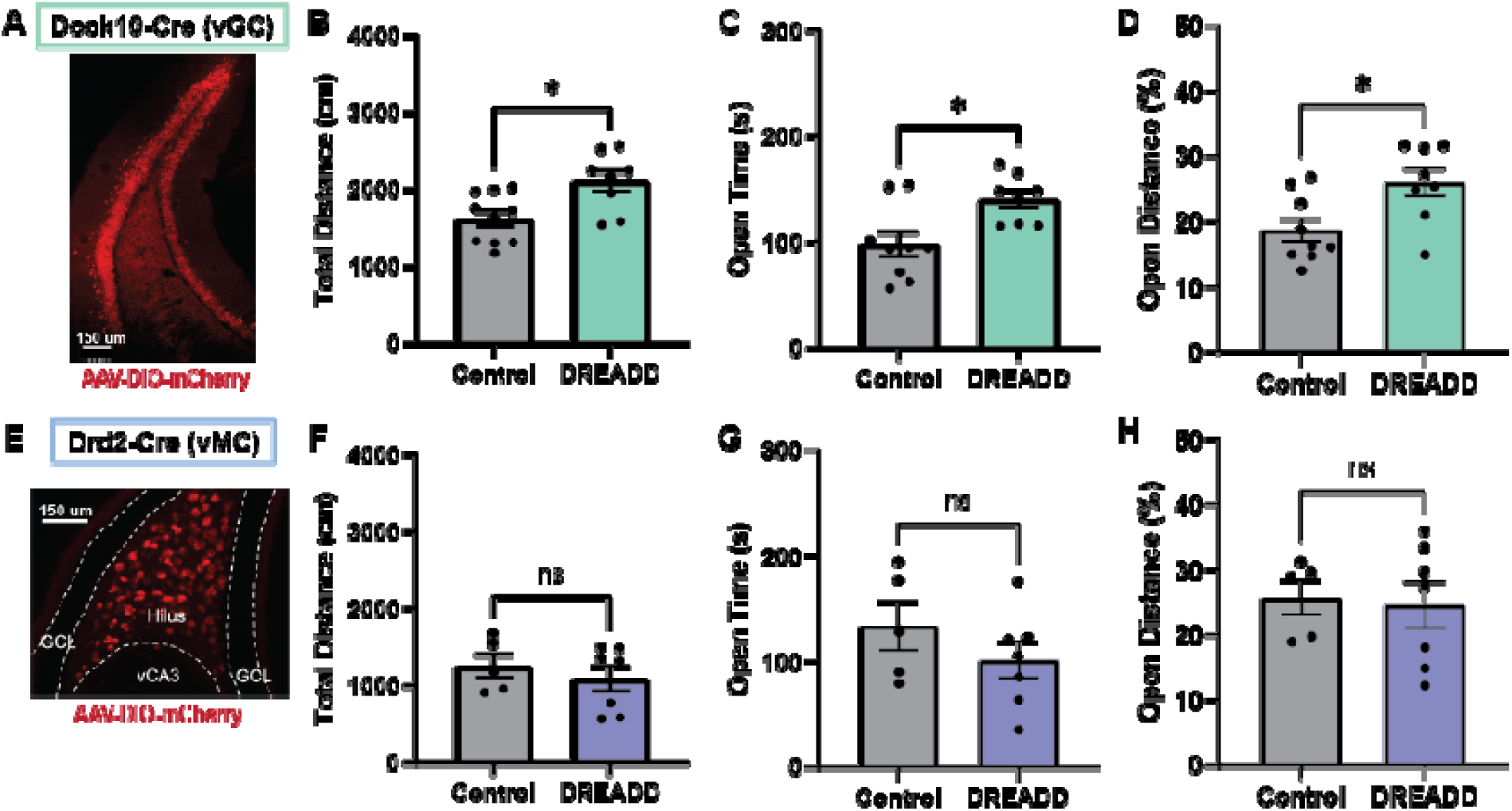
vGCs, but not vMCs affect innate avoidance in the elevated plus maze. AAV-DIO-mCherry (Control) or AAV-DIO-hM4Di (DREADD) was injected into the vDG of Dock10-Cre (A-D) or Drd2-Cre (vMC) male mice. (A) AAV-DIO-mCherry expression is limited to vGCs in Dock10-Cre mice. (B-D) Silencing vGCs while mice explore the EPM increased total distance traveled (B), time spent in open arms (C), and percent distance traveled in the open arms (D), indicating decreased avoidance behavior. AAV-DIO-mCherry expression is limited to vMCs in Drd2-Cre mice. (F-H) Silencing vMCs while male mice explore the EPM did not significantly affect total distance traveled (F), time spent in open arms (G), or percent distance traveled in open arms (H). Error bars represent SEM. * p<0.05. N_control_ = 10; N_DREADD_ = 8.

## DISCUSSION

The ventral hippocampus is critical for emotional behavior, including approach-avoidance conflict resolution in rodents^4,14,15^. Lesion and circuit manipulation studies have implicated both vDG and vCA1 in such behaviors, however the neural population activity that supports this behavior is not well understood. In this study, we used transgenic mouse lines to selectively record or manipulate activity in mossy cells or granule cells in the ventral dentate gyrus, as well as vCA1, during exploration of a novel elevated plus maze and open field in order to test how these different populations encode specific aspects of anxiety-like behavior.

Previous studies have found that vCA1, but not dCA1, neurons have a rate preference for the anxiogenic open arms of the EPM^5^, indicating a specific relationship of the ventral hippocampus to anxiety-like features of an environment. Furthermore, vCA1 and vDG are essential brain structures for normal behavior in approach-avoidance tasks such as the EPM and NSF^3,15–17^. However, specific coding properties of neurons in vDG have not been investigated in an anxiety-related task. It is therefore unknown whether anxiety-related information is encoded similarly in vDG and vCA1, and thus whether it is computed directly by the CA1 circuit, or emerges by virtue of inputs from upstream hippocampal subregions such as the dentate gyrus. vCA1 has extensive bidirectional connections with the basolateral amygdala^18,19^, and the BLA-vCA1 projection has been implicated in the expression of normal avoidance behavior in the EPM^20^. Therefore, it is possible that the anxiety-related firing identified in vCA1 originates entirely from BLA and not from within the trisynaptic circuit itself.

However, we demonstrate that both populations of vDG neurons, vGCs and vMCs, have a strong firing preference for the open arms of the EPM, which suggests that this feature of vCA1 neurons is present at the first stage of information processing in the ventral hippocampus, the dentate gyrus. Furthermore, all three cell types exhibited increased event rates during risk assessment behaviors such as head dipping and stretching, which were also previously observed in vCA1.

Interestingly, whereas vGCs and vCA1 exhibited similar distributions of neurons that preferred each arm type, closed or open, a significantly greater proportion of vMCs had an open arm preference (**Figure 2**). Neural activity may increase in the open arms for a variety of reasons, including increased visual flow, general arousal, novelty, and negative valence. The fact that vMCs were particularly sensitive to open arm exploration and risk assessment (**Figure 2A**), yet did not encode arm type in an abstract manner (**Figure 4F-H**), indicates that the open arm rate increases seen in vHPC may not be an accurate indicator of generalized anxiety encoding. Or, it may be that the higher spatial encoding of vMCs precludes abstraction between two spatially distinct arms. vMCs also appear to be particularly sensitive to novelty and exploratory behaviors. In the EPM, they demonstrate a rapid decrease in activity within the first two minutes of the task, whereas vGCs and vCA1 do not return to similar relative rates until the end of the session (**Figure 2A**). In addition, rearing against the walls in an open field elicits a significantly greater increase in activity in vMCs compared with the other cell types (**Figure S4**). Rearing is considered an exploratory behavior in rodents, as it enables the animal to gain a different perspective of their surroundings^21^, which may be consistent with the novelty responses of vMCs in other measures. Furthermore, while all cell populations exhibited decreased overall activity in the closed arms of the EPM relative to open arms, vMCs were unusual in that they responded briefly with an increase in activity upon entering a closed arm (**Figure 2E**). Thus, vMCs appear to be more activated by changes in the environment than either vGCs or vCA1. This is consistent with recent work in which vMC calcium activity decreased during multiple exposures to a familiar open field environment^22^. Such state-dependent responses may be due to actions of one of the many neuromodulators that selectively target mossy cells, cell intrinsic properties, or distinct inputs from the diverse subtypes of interneurons in the region.

The hippocampus has traditionally been considered to be critical for spatial navigation-neurons in all dorsal hippocampal subfields have been found to convey spatial information, and the existence of place cells in each subregion has been well documented^11,23–25^. It has also been observed that there is a functional gradient along the longitudinal axis, in which dorsal, but not ventral HPC lesions lead to spatial memory deficits^26,27^, and the spatial scale of hippocampal place fields is larger in ventral hippocampus compared with dorsal^28^. Recently, population-level decoding analyses have expanded our understanding of the mechanism by which dorsal hippocampal ensembles encode spatial features such as position, velocity, and head direction^10^. We now show that the animal’s location in two different types of arenas can also be decoded from ventral hippocampal populations as well.

In this study, we first quantified the spatial information conveyed by individual vGCs, vMCs, and vCA1 neurons and found that while vGCs and vCA1 neurons carry a relatively low amount of spatial information, vMCs encode spatial information with significantly higher precision in both EPM and open field environments. <u>Next, in order to</u> explore whether spatial position is encoded in vHPC ensembles at the population level, we used a linear decoder to predict the location of the mouse in both the EPM and open field tests. In both arenas, position could be decoded better than chance from all vHPC cell types. However, vMCs provided significantly better position decoding performance than the other cell types, which is consistent with the single-cell information metric.

Thus, there is a dissociation within vDG such that vGCs convey lower spatial information similar to vCA1, and vMCs convey higher spatial information similar to dorsal HPC. The origin of this spatial information, however, remains unclear, given that the majority of vMC inputs arise from vGCs, local interneurons, and neuromodulators such as serotonin and dopamine. It is possible that vMCs receive spatial information from dorsal CA3^29^, or dorsal mossy cells which are known to innervate the ventral DG^30^, or alternately emerge via combinations of inputs from vGCs with weaker spatial coding. Spatial information from vMCs may subsequently feed back upon vGCs in order to contribute to the spatial context of ventral hippocampus approach/avoidance representations.

To further examine ventral hippocampus anxiety representation at the population level, we implemented a decoding strategy that has been used to assess abstract population coding in human and nonhuman primates: multidimensional scaling (see Methods)^13,31^. This decoding strategy showed that vGCs and vCA1 neurons encode arm type in an abstract sense such that arm valence could be generalized to new arms, whereas vMCs did not. Taken together, the picture that emerges is one in which vGCs and vCA1 share many of the same properties that enable these neurons to process approach-avoidance conflict information.

The implication of these findings is that anxiety-related information is present even at the initial stage of hippocampal processing, the dentate gyrus. However within the DG there is a segregation between the two excitatory cell types, with vGCs being more tuned to anxiety and vMCs to spatial position. To test whether vGC activity is important for anxiety-related behavior, we chemogenetically inhibited vGCs or vMCs while mice explored the EPM. As predicted, silencing vGCs, but not vMCs, resulted in increased exploration of the open arms, an indication of decreased anxiety. Though vMCs exhibited increased open arm activity, they did not encode arm type but rather spatial position and novelty. Interestingly, MCs project broadly within the DG both along the dorso-ventral axis and contralaterally. It is therefore possible that they receive their spatial information from the dorsal hippocampus and in turn broadcast it to the whole hippocampus. Thus, different features of a novel, anxiogenic context are differentially encoded within different neural populations of the ventral hippocampus. It remains to be determined which inputs to the ventral hippocampus are necessary to enable the formation of valence-related representations within the trisynaptic circuit.

## METHODS

### Animal Subjects

Procedures were conducted in accordance with the U.S. NIH Guide for the Care and Use of Laboratory Animals and approved by the Institutional Animal Care and Use Committees of the New York State Psychiatric Institute. Male and female Dock10-Cre and Drd2-Cre transgenic mice were bred on a C57/Bl6 background, and were graciously provided by Susumu Tonegawa and Helen Scharfman, respectively. Data from male C57/Bl6 mice used for vCA1 calcium imaging recording were generated in a previous study from our lab^5^. Mice were utilized for experiments at 8-14 weeks of age and given unrestricted access to food and water on a 12-hr light cycle. Experiments were performed during the light portion of the cycle.

### Viral Constructs

Adeno-associated viruses for calcium imaging were supplied by AddGene or UPenn Vector Core. AAV1-Syn-GCaMP6f.WPRE.SV40 was used to label neurons in vCA1, and was provided by UPenn Vector Core at a titer of ∼6 × 10^12^ vg/mL and diluted to ∼2 × 10^12^ vg/mL prior to injection. AAV5-Syn-Flex-GCaMP6f.WPRE.SV40 was used to label mossy cells in the ventral dentate gyrus, and was provided by Penn Vector Core at a titer of ∼4 × 10^12^ vg/mL. AAV9-Syn-FLEX-jGCaMP7s-WPRE was provided by AddGene at a titer of ∼1 × 10^13^ vg/mL and diluted to ∼3 × 10^12^ for injection into Dock10-Cre mice to label dentate granule cells.

### Stereotactic Surgeries

During all surgical procedures, mice were anesthetized (1.5% isoflurane, 1 L/min O_2_) and head-fixed in a stereotaxic frame (David Kopf, Tujunga, CA). Fur near the incision site was shaved and the region sterilized with betadine and 70% ethanol, and ophthalmic ointment was applied for eye lubrication. Body temperature was maintained with a T/pump warm water recirculator (Stryker, Kalamazoo, MI). Subcutaneous saline and carprofen was given peri-operatively and post-operatively for 3 days for hydration and analgesia. *In vivo* Ca^2+^ imaging surgical procedures were conducted as previously described ^5,32^. Briefly, 3 small screws (FST, Foster City, CA) were inserted into burr holes around the implantation site, where a craniotomy was made and dura removed from the brain with fine forceps. Mice were injected unilaterally with virus. For vCA1 mice, mice were injected (-3.16 AP, 3.25 ML, -3.85, -3.5, 3.25 DV measured in mm from the brain surface) with AAV-syn-GCaMP6f with a Nanoject syringe (Drummond Scientific, Broomall, PA) followed by implantation of a 6.1mm long, 0.5mm diameter GRIN lens (Inscopix, Palo Alto, CA). The lens was lowered slowly in 0.1mm DV steps to the target site (vCA1: -3.16 AP, 3.5 ML, -3.5 DV in mm from skull) and fixed to the skull with dental cement (Dentsply Sinora, Philadelphia, PA). For vMC and vGC mice, a similar procedure was used with the exception that the lens was implanted using the ProView system (Inscopix, Palo Alto, CA) 3 weeks following viral injection, in order to visualize fluorescence during lens implant and thus increase the surgery success rate. For vMC, AAV5-Syn-Flex-GCaMP6f was injected unilaterally at 2.8 ML, -3.6 AP, -3.5 DV mm as measured from bregma. For vGC mice, AAV9-Syn-Flex-GCaMP7s was injected unilaterally at the same coordinates. The target depth for vMCs and vGCs was -3.5mm and -3.1mm DV as measured from the skull surface, respectively.

### Behavioral Assays

#### Elevated Plus Maze

Mice were placed in a standard-sized EPM (34 cm height from floor, 63.5 cm full arm length, 5.5 cm arm width, 17.8 cm tall closed arms, with 1cm wide ledges on the open arms). Lamps were placed to achieve ∼300-600 lux centered over the open arms to promote avoidance behavior. Mice were placed in the center of the maze facing a closed arm, and were allowed to explore for 10 minutes while recording behavior with a C920 webcam (Logitech, Lausanne, Switzerland) or Basler ace GigE monochrome digital camera (Basler, Ahrensburg, Germany).

#### Open Field Test

Mice were placed in an open field arena (44 cm x 44 cm x 30 cm LWH; Kinder Scientific, Poway, CA) with a bright light ∼400-600 lux centered over the center of the maze. Mice were allowed to explore for 10 minutes while behavior was recorded with a C920 webcam (Logitech, Lausanne, Switzerland) or Basler ace GigE monochrome digital camera (Basler, Ahrensburg, Germany).

### Pose Estimation

Behavior videos were analyzed using the DeepLabCut software package ^33^. A neural network was trained to identify six mouse body parts: nose, both ears, head, body, and tail base. These body parts were manually labelled on a set of ∼30-100 still frames from each behavior video, and the network was trained to detect these body parts. After the neural network was sufficiently trained (train root mean square error < 4 pixels), the tracking data was further processed using custom python code. For each behavior video, a single frame was extracted and regions of interest (ROIs) were drawn over the image, and the coordinates were saved. In the EPM, the ROIs drawn were: arena (a very loose outline of the entire maze), open arms, and closed arms. In the open field, the arena and floor were drawn. Behavior zones (e.g. open/closed arms in the EPM, floor in OFT) were carefully drawn such that they accurately identified zone boundaries. Next, tracking data was processed to reduce noise and errors. First, frames in which a body part was located outside of the arena or were predicted with low likelihood (< 0.9) were deleted, and the time series was interpolated to fill in missing data. Next, the data was smoothed using a Savitzky-Golay filter, and boolean time series were generated for each body part and ROI to indicate whether the body part was located within each ROI at a given time point.

Beyond the (x,y) location of each body part, features of the animals’ pose were extracted from the tracking data. Egocentric and allocentric head direction were computed using the vector joining the left and right ear, and body length and orientation were computed using the vector joining head, body, and tail base. From some of these features, individual behaviors were defined and validated with manual scoring. Head dips were defined as time points in which the head was located in the open arms and was within 1cm of the edge of the open arm zone. Stretching was defined as time points during which the body length of the mouse exceeded a threshold that was determined for each mouse. The body length threshold was empirically determined to be: 1.2*c, where c is the median body length when the body point velocity is between 1 cm/s and 10 cm/s. The velocity was restricted in this way to eliminate time points when the mouse was moving quickly or was stationary, as the measured body length during these time periods is not reflective of the true body length. Rearing in the open field was defined by time points in which the head was located within 1cm of the edge of the floor, as this captures moments when the mouse is rearing against the wall of the arena. Unfortunately, a reliable method for detecting rears in the middle of the arena was not discovered, though the majority of rearing events occurred against the wall.

### Freely Moving Calcium Imaging

vGC and vMC mice underwent a similar imaging protocol to vCA1 mice, as previously described ^5^. Three weeks following lens implant surgery, mice were head-fixed while anesthetized with 1.5% isoflurane with 1 L/min oxygen in order to check GCaMP expression with the miniature microscope (Inscopix, Palo Alto, CA). The protective rubber mold was removed from the lens, and the lens was cleaned with lens paper. A baseplate was screwed onto the microscope and lowered over the GRIN lens until the field of view came into focus. If fluorescent cells could be visualized, dental cement was applied to the baseplate to fix it to the headcap. After the cement hardened, the baseplate was unscrewed from the microscope, which was then removed and replaced with a lens cover (Inscopix, Palo Alto, CA). Prior to each imaging session, mice were anesthetized briefly for ∼2 min to attach the miniscope to the baseplate, after which mice were allowed to recover for 30 min before beginning imaging. Calcium videos were recorded with nVista acquisition software (Inscopix, Palo Alto, CA), and behavior and calcium imaging videos were synchronized with a TTL-triggering system (a TTL pulse from Ethovision XT11 and Noldus IO box was received by nVista DAQ box to begin imaging session). Calcium videos were acquired at 20 frames per second and the same LED power settings were used for each mouse.

### Histology and Confocal Microscopy

After completing the experiment, mice were anesthetized and transcardially perfused with 4% paraformaldehyde (PFA) in 1X phosphate buffer solution (PBS), after which brains were fixed in 4% PFA overnight. Brains were transferred to 30% sucrose for 2 days, after which they were frozen in methylbutane and coronally sliced on a cryostat (Leica CM 3050S) at a thickness of 50 um. Sections were mounted and cover slipped with Vectashield antifade mounting medium with DAPI (Vector Laboratories, Burlingame, CA).

### Image Processing

All raw calcium videos, including the previously-recorded vCA1 videos, were processed in the same manner using Inscopix Data Processing software (versions 1.1 and 1.2, Inscopix, Palo Alto, CA). Videos were temporally downsampled to 10 frames per second, spatially downsampled by a binning factor of 4, and motion corrected. Neurons were segmented using Constrained Non-negative Matrix Factorization for microEndoscopic data (CNMF-E)^34^. Identified neurons were sorted by manual inspection for appropriate spatial footprints and calcium temporal dynamics. Three temporal signals were considered as the output of CNMFE: the raw calcium trace (C_raw), the deconvolved trace (C), and the deconvolved spikes (S), which provides an estimate of the number of spikes that occurred at each time bin. A minimum spike size (s_min) of 10 * estimated noise (sn) was applied to eliminate numerous small-amplitude spikes estimated from the calcium trace. All three traces were normalized by the standard deviation of the noise (defined as C_raw – C).

### Quantification and Statistical Analysis

#### Calcium Data Analysis

All data analysis was performed using the Caliban python software package (www.github.com/jaberry/caliban), which provides a suite of modules for performing common single-cell and population level analyses. In this paper, calcium rate was calculated as the total number of normalized estimated spikes divided by the time. This metric differs from previous work that has used event rate to compare neural activity, in which each major calcium transient is assigned an event size of 1. Event rates have a few major disadvantages compared with using normalized estimated spike rates. First, each transient is treated equivalently in the rate calculation, and thus does not capture the variability in transient size. Second, many algorithms for detecting transients rely on a single amplitude threshold that can be arbitrarily set and are therefore not standard across experiments. Finally, transient detection using simple cutoffs struggle to accurately detect multipeaked transients, which are quite common in highly active cell types such as mossy cells^35^. Unless otherwise stated, the normalized deconvolved spike trace was used for calcium activity analysis.

#### Task-selectivity Analysis

Neurons were classified as selective for a particular behavioral variable (e.g. open arm exploration) as previously described^5^. Briefly, deconvolved spikes were shuffled 1000 times and the mean rate was computed for time spent during the behavior, and not during the behavior, and the difference mean rates was recorded. The true (non-shuffled) mean rate difference was compared to the shuffle distribution using a nonparametric two-sided permutation test. A neuron was deemed selective for that behavior if the permutation test p-value was less than 0.05, and if the rate difference was greater than 0 (a negative rate difference would indicate the neuron was significantly inhibited during the behavior). A neuron that was not selective for any behavior was considered ‘not-selective’.

#### Spatial Information

The spatial information content carried in the activity of each cell was computed as previously described^10,36^. We divided the entire arena into 4×4 cm squares and computed the Shannon information that a single event conveys about an animal’s location. The spatial information carried by each cell was computed as the mutual information score between event occurrence and animal location. To correct for sampling bias in information measures^37^, we shuffled the neural time series 1000 times and computed the spatial mutual information for each shuffle. The significance of spatial information (SSI) was computed as the number of standard deviations away from the mean of the shuffle distribution.

#### Position Decoding

Position was decoded in both EPM and OFT using a Support Vector Machine (SVM) classifier with a linear kernel, as previously described^10^. The arena was divided into a 16×16 grid of 64 discrete locations, and a SVC was used to predict the location of the mouse. Decoder performance was assessed using 10-fold cross-validation, in which the data was divided into 10 groups and one group was held out as the test set. Decoder error was assessed as the distance in cm between the predicted location and the actual location. To control for variability in calcium decay among cells, an exponential decay function with fixed tau was convolved with the deconvolved spikes to generate a novel deconvolved trace. In this paper, a decay constant of tau = 2s was used for all decoding.

In order to determine the statistical significance of the decoder performance, we generated a chance distribution of decoding errors. This was achieved by reversing and shifting in time the position vector by a random interval, and testing decoder performance on this shifted data. The real performance was then compared with the shuffled performance distribution using a Mann-Whitney statistical test.

#### EPM Score

The EPM score was computed as previously described^6,12,38^ as a single cell measure of anxiety-related neural activity. The EPM score represents a ratio of the mean difference in activity between arms of the same type with arms of different type. It ranges from - 0.33 to +1, with positive values corresponding to neurons that are active in both arms of the same type, but not in either of the other arms. Scores near zero correspond to neurons that do not have an arm-type preference. Negative scores indicate activity that is similar among arms of different type. The formula used is:

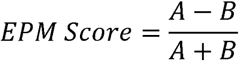

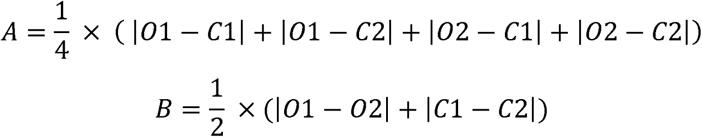

Here, O1, C1, O2, C2 refer to the normalized calcium rate in each arm (open1, closed1, open2, closed2. The quantity A represents the mean rate difference between arms of different valence, and the quantity B represents the mean rate difference between arms of the same valence.

#### Decoding test for arm generalization

We used a decoding strategy to assess whether the neural population encodes information in a manner that allows for information to be generalized within classes of related behavioral variables. This strategy has previously been used to assess whether variables are encoded in an abstract manner in primates and human HPC and PFC^13,39^. A traditional k-fold cross validation decoding strategy would involve training a decoder on 90% of the data and testing on the remaining 10%. While this strategy quantifies the ability of a neural population to encode each variable or context, it cannot inform us as to whether the variables are encoded in an abstract manner, which would allow for generalization to new variables that share features with the originally encoded variables ^13^. To assess whether anxiety-related information is stored in such an abstract manner in vHPC, the data was split between arms of the same type. For example, Open1 and Closed1 may be paired together in one group, and Open2 and Closed2 in the other group. Alternatively, pairs could be Open1/Closed2, and Open2/Closed1. An SVC with a linear decoder was trained to distinguish a variable such as arm type (Open vs Closed) for EPM, or Center vs Corners for OFT, using a single pair. The performance of the decoder was evaluated on the remaining pair, thus testing whether the decoder was able to generalize that variable identity to a novel context.

Both the EPM and OFT are anxiety-related tasks that by design comprise ‘safe’ and ‘unsafe’ zones. Thus, these zones will have unequal occupancy, which can bias traditional classification performance metrics such as accuracy, precision, and recall. To overcome this bias, we used the Matthews Correlation Coefficient (MCC)^40^, which is a balanced measure of binary classification performance^41^. The formula for MCC is as follows:

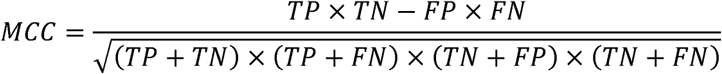

Where TP, TN, FP, and FN indicate the number of true positives, true negatives, false positives, and false negatives, respectively. This MCC metric ranges from -1 to +1, with 0 indicating chance levels of classification.

To control for the variability in the number of cells in each FOV, 20 cells were randomly selected from each mouse for decoding. The results from 25 iterations was aggregated to estimate the decoding performance for each mouse. Additionally, the significance of the performance was estimated by iteratively shifting the behavior vector by random intervals and repeating the decoding procedure to generate a chance distribution. The true result was compared to the shuffle distribution using a Mann-Whitney test, and p-values were reported as -log(p) for each cell group.

#### Multidimensional Scaling Plots

The geometry of the neural representations of the EPM were visualized using multidimensional scaling^13,31^. A population vector representing the average event rate for each neuron was computed for each entry into an arm that lasted longer than 0.5 seconds. The population vectors were normalized, and MDS was computed using the scikit-learn MDS function. This was used to map the population activity during each arm entry onto 2-D space. This technique enables us to understand the relationship between individual arm representations in different neural populations.

## ACKNOWLEDGEMENTS

The authors would like to thank Sean Lim, Kassandra Carrion, Gergely Turi, Fabio Stefanini, Helen Scharfman, Steven Siegelbaum, and Attila Losonczy for technical and scientific discussions. This work was funded and supported by the National Institutes of Health (Grant Nos. F30MH1178927 [Principal Investigator (PI): JEB], R37MH068542 (JEB, JCJ), R25MH06048 (JEB), K08MH122893 [PI: WC], R01MH068542 [PI: RH], RF1AG080818 [Co-PI: RH]) Funding and support also provided by the Hope for Depression Research Foundation [PI: RH, Co-Investigator WC].

