## Supplemental Figures for "Distinct representations of an anxiogenic environment in different cell types of the ventral hippocampus"

**Figure S1.**

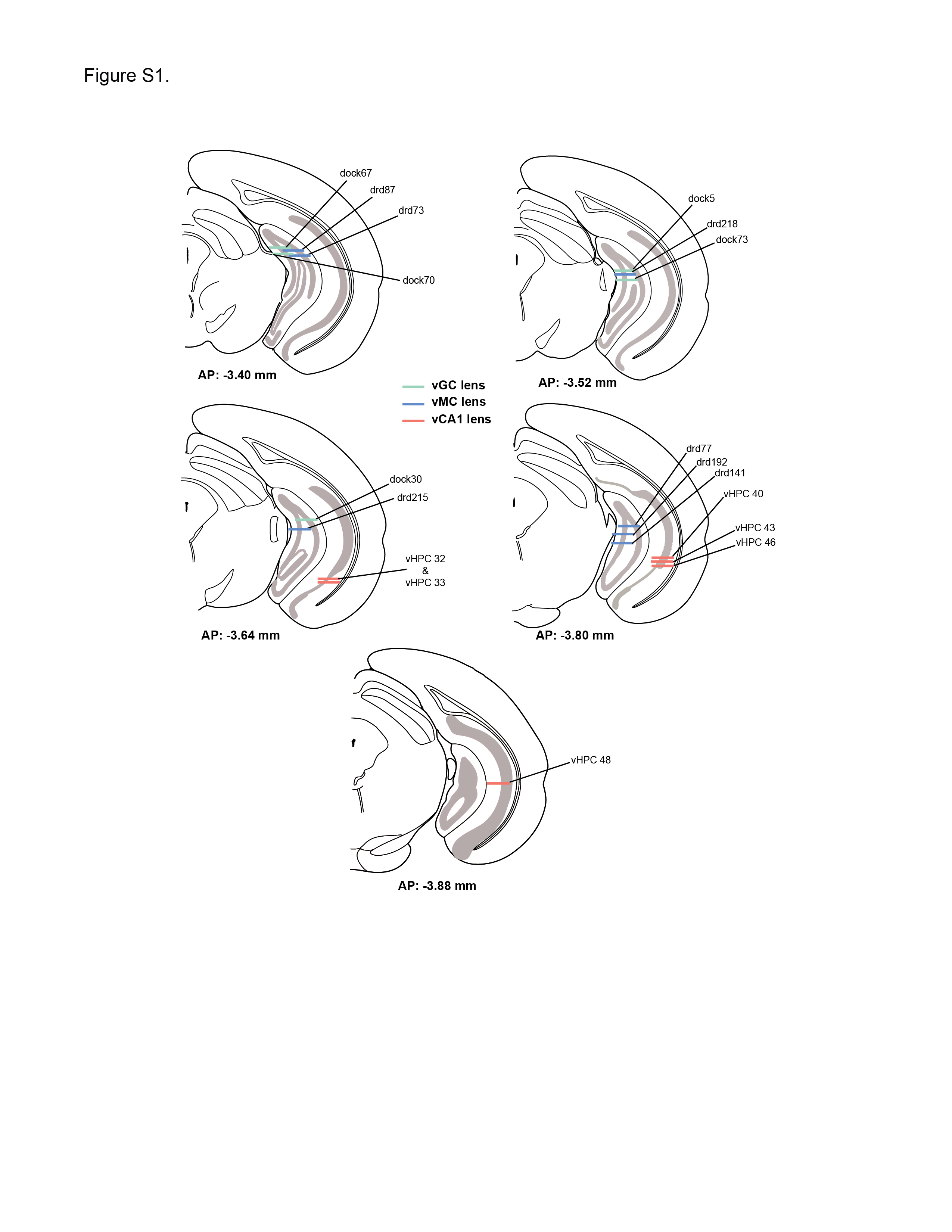

Figure S1. Lens placements for each mouse. The location of the bottom of the GRIN lens for each mouse in this study. Dock10-Cre mice used to image vGCs are shown in green. Drd2-Cre mice used to image vMCs are shown in blue. C57/Bl6 mice used to record from vCA1 cells are depicted in orange.

**Figure S2.**

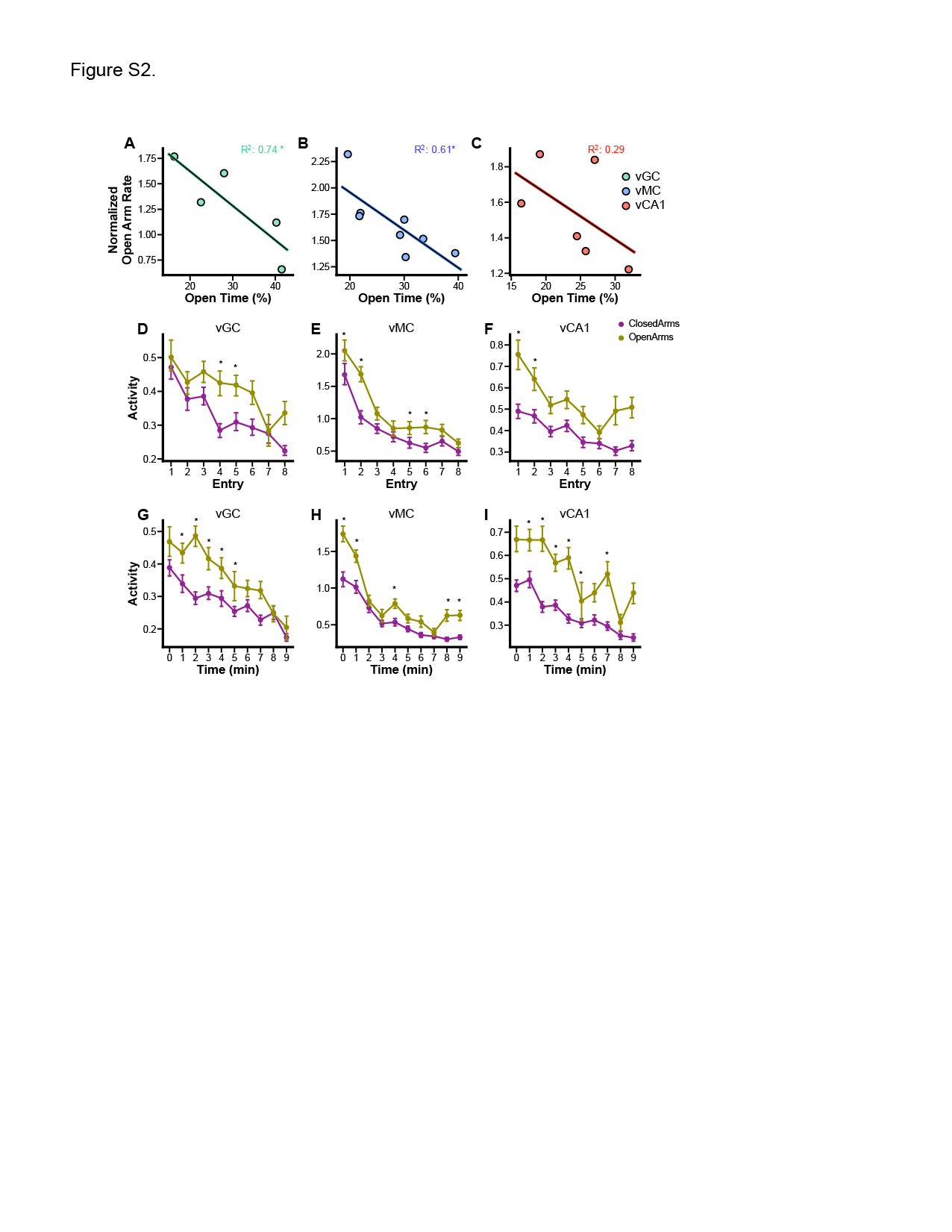

Figure S2. Open arm calcium event rate correlates inversely with anxiety level, and open arm rate preference persists across time in the EPM**.** (A-C) Mean normalized open arm rate is correlated with time spent in the open arms for each cell type. Normalized rate are calculated for each neuron by dividing the rate in the open arms by the mean rate during the entire session. (A) vGCs: R^2^ = 0.74, p = 0.06. (B) vMCs: R^2^ = 0.61, p = 0.02. (C) vCA1: R^2^ = 0.29, p = 0.27. (D-F) Mean rates in open and closed arms decline over the first 8 entries into each arm type, but the open arm rate preference is largely preserved. (G-I) Similarly, average rates decreased over the 10 minutes of EPM exploration in all cell types.

**Figure S3.**

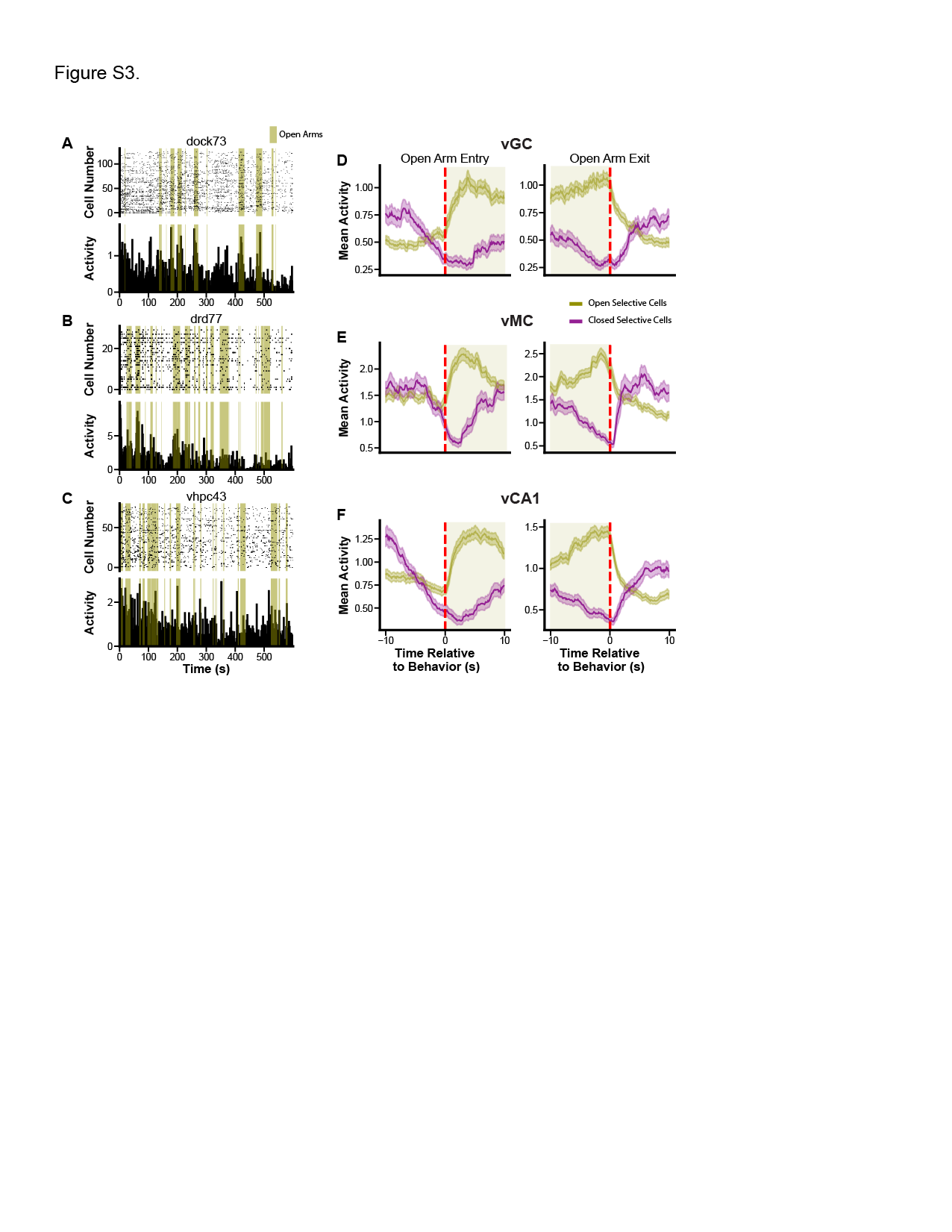

Figure S3. Population level open arm rate preference derives from subpopulations of open arm-selective cells. (A-C) Raster plots from representative mice for vGC (A), vMC (B), and vCA1 (C) for the entire 10-minute EPM session. *Top*: raster plot. *Bottom*: Average rate per 2s time bin. Green bars indicate time spent exploring the open arms. (D-F) Event triggered average calcium plots for open arm-selective cells (green) and closed arm-selective cells (purple) for vGC (D), vMC (E), and vCA1 (F) cells. Left: Activity relative to entering an open arm. Right: Activity relative to exiting a closed arm. Each cell group is composed of neurons that are preferentially active in one arm type. Note that both open/closed arm selective cell rates remain elevated/decreased after initial open arm entry, respectively, for vDG and vCA1, however vMCs rapidly return to baseline levels, consistent with a response to novelty.

**Figure S4.**

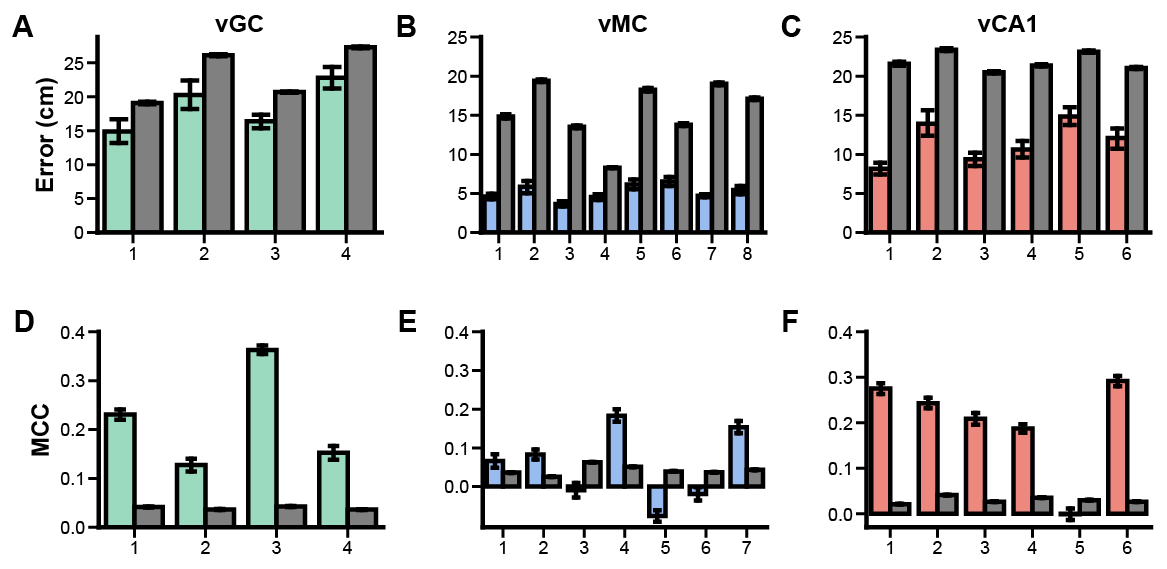

Figure S4. Decoding position and arm type in the EPM for each mouse**. (**A-C) Position decoding performance for individual mice from vGC (A), vMC (B) and vCA1 (C) groups. Grey bars, chance performance distribution. Color bars: actual performance. (D-E) Arm type decoding performance for each mouse when decoding from vGCs (D), vMCs (E), and vCA1 (F). Grey, chance performance; Color; True performance. Performance metric is Matthews correlation coefficient (MCC).

**Figure S5.**

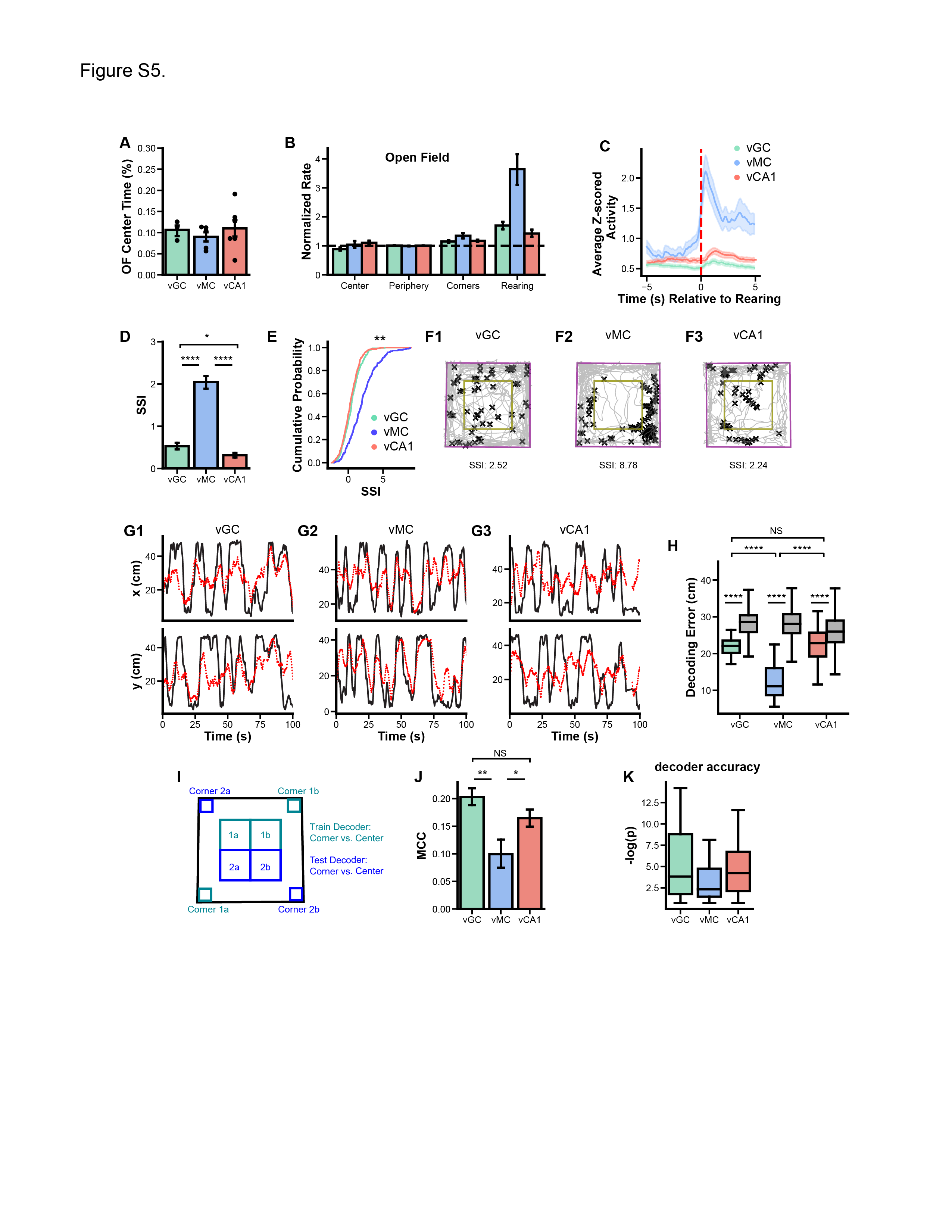

Figure S5. Ventral hippocampal activity in the Open Field Test. (A) Time spent in the center of the open field was not significantly different among mice in which different cell groups were recorded. (B) Normalized calcium event rates in the center, periphery, and corners of the open field, and during rearing events. Normalized rate for each neuron is calculated by dividing the rate during these epochs by the mean rate during the entire session. Rates were not significantly elevated during center exploration, however they were elevated during rearing. In particular, vMCs were highly activated during rearing. (C) Rearing-triggered average of mean rates. (D-E) Mean standardized spatial information (SSI) is significantly higher in vMCs compared with vGC or vCA1. (F-H) Representative activity plots for single vGC (F1), vMC (F2), and vCA1 (F3) neurons. (G-H) Spatial decoding in the open field. (G) Open field actual position (black) and decoded position (red) for one open field session from vGC (G1), vMC (G2), and vCA1 (G3) mice. (H) Mean spatial decoding error in each cell type vs. shuffled data (gray). (I-K) The ability of the population to generalize across locations with similar valences was tested in a manner similar to that used in EPM. (I) Illustration of the decoding strategy used. A linear decoder was trained to distinguish whether the mouse was in the center or corners using half of the data (blue regions), and tested on the remaining half (teal regions). (J) The performance of the decoder, as measured by the Matthews correlation coefficient (MCC). Center vs. corner region decoding performance was worse when using vMC activity compared with vGC or vCA1. (K) Mann-Whitney test p-values, expressed as –log(p), decoder performance using real vs. shuffled data.

**Figure S6.**

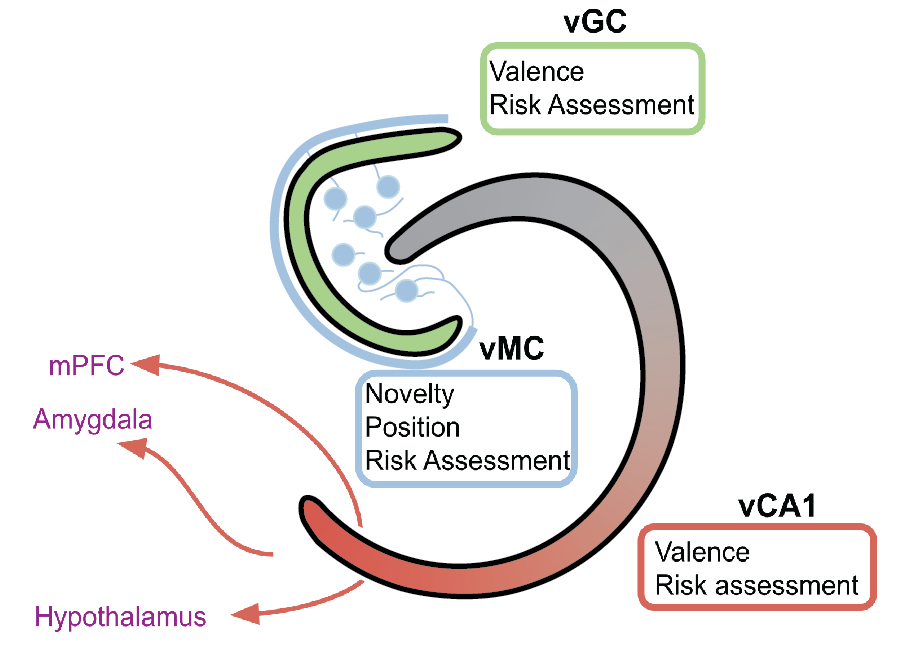

**Figure S6. Model of spatial and anxiety-related encoding within the ventral hippocampus**. vGCs (green) and vCA1 (red) encode abstract information about EPM arm type or valence, whereas vMCs respond to novelty and encode spatial position. All three cell types are active during risk assessment behavior such as open arm exploration, head dips, and stretching.
